# Coalescent theory of migration network motifs

**DOI:** 10.1101/190249

**Authors:** Nicolas Alcala, Amy Goldberg, Uma Ramakrishnan, Noah A. Rosenberg

**Affiliations:** Department of Biology, Stanford University, Stanford, CA 94305-5020, USA; National Centre for Biological Sciences, Bangalore, Karnataka 560065, India

**Keywords:** coalescent theory, genetic differentiation, network, population structure

## Abstract

Natural populations display a variety of spatial arrangements, each potentially with a distinctive impact on genetic diversity and genetic differentiation among subpopulations. Although the spatial arrangement of populations can lead to intricate migration networks, theoretical developments have focused mainly on a small subset of such networks, emphasizing the island-migration and stepping-stone models. In this study, we investigate all small network motifs: the set of all possible migration networks among populations subdivided into at most four subpopulations. For each motif, we use coalescent theory to derive expectations for three quantities that describe genetic variation: nucleotide diversity, *F*_*ST*_, and half-time to equilibrium diversity. We describe the impact of network properties on these quantities, finding that motifs with a large mean node degree have the largest nucleotide diversity and the longest time to equilibrium, whereas motifs with small density have the largest *F*_*ST*_. In addition, we show that the motifs whose pattern of variation is most strongly influenced by loss of a connection or a subpopulation are those that can be split easily into several disconnected components. We illustrate our results using two example datasets—sky island birds of genus *Brachypteryx* and Indian tigers—identifying disturbance scenarios that produce the greatest reduction in genetic diversity; for tigers, we also compare the benefits of two assisted gene flow scenarios. Our results have consequences for understanding the effect of geography on genetic diversity and for designing strategies to alter population migration networks to maximize genetic variation in the context of conservation of endangered species.

---

COALESCENT theory is a powerful tool to predict patterns of genetic variation in models of population structure, and many studies have investigated the predictions of coalescent models about genetic variation under a variety of different assumptions about the genetic structure of populations (Donnelly and Tavaré 1995; Fu and Li 1999; Rosenberg and Nordborg 2002).

Correctly predicting the effect of connectivity patterns on the expected amount of nucleotide diversity and genetic differentiation is important in a range of settings. In population genetics, such predictions enable descriptions of the impact of migration as one of the main evolutionary forces influencing allele frequencies. In molecular ecology, they help evaluate the consequences of abiotic factors such as geographic barriers, and biotic factors such as assortative mating, on levels of genetic diversity and genetic differentiation. In conservation genetics, they can be used to quantify the impact of past and future disturbance, as well as to predict the outcome of management initiatives.

The two most frequently examined models of population structure are the island-migration and stepping-stone models. In the island model, individuals can migrate from any subpopulation to any other subpopulation, all with the same rate (Wright 1951). In the stepping-stone model, individuals can only migrate to neighboring subpopulations (Kimura 1953; Maruyama 1970). Stepping-stone models can represent multiple spatial arrangements. Under the circular stepping-stone model, subpopulations are arranged in a circle, so that all individuals can migrate to exactly two subpopulations.

Although the island and stepping-stone models can accommodate a variety of patterns of connectivity among subpopulations, they represent only some of the possible patterns, or network “motifs.” Indeed, these models account for only 7 of 18 motifs possible for sets of one to four subpopulations (Figure 1). Numbering motifs by the classification from Read and Wilson (2005, p. 8), motif 1 corresponds to the panmictic population model, motif 18 to the island model, motifs 6,14, and 16 to stepping-stone models, and motifs 3 and 7 to both island and stepping-stone models. Although tools of coalescent theory to study arbitrary migration models are available (Wilkinson-Herbots 1998), to our knowledge, patterns of variation expected from the remaining 11 motifs have not been described.

**Figure 1.**
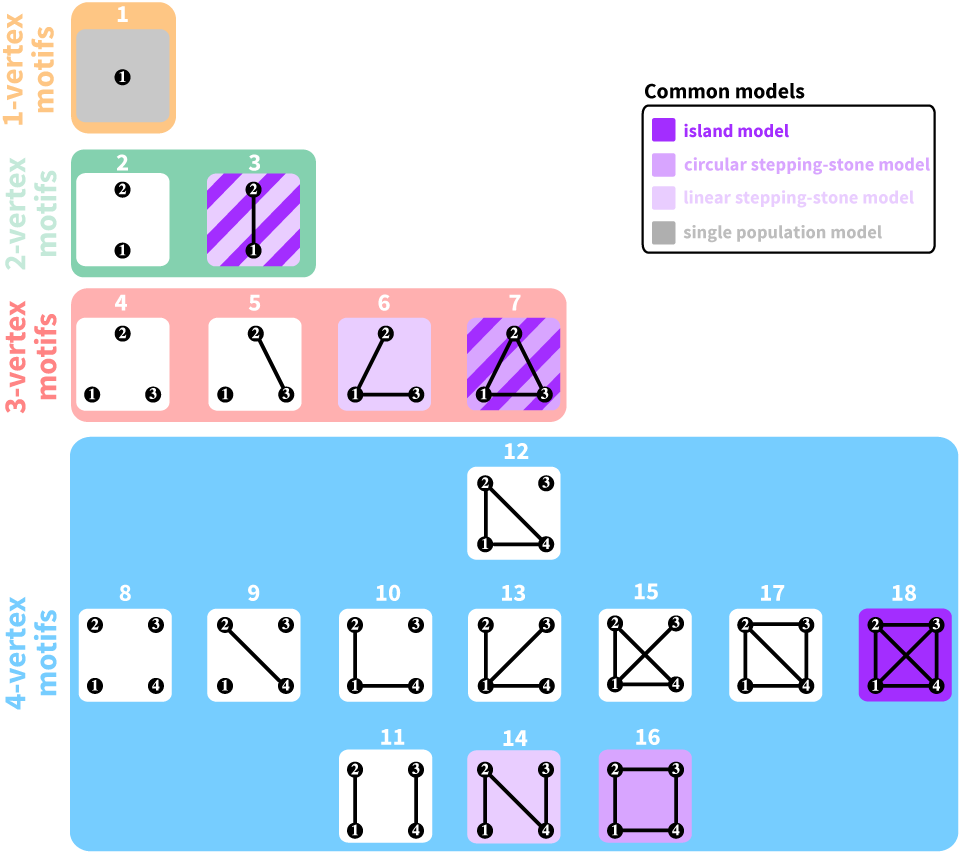
All possible network motifs for sets of at most 4 vertices. Purple motif backgrounds highlight motifs that follow standard models, island or stepping-stone or both. Note that we take the term "motif" to indicate a specific small undirected graph (rather than a small directed or undirected subgraph statistically overrepresented in large empirical networks, as in many applications).

An objective in the study of spatial arrangements of populations is to examine the properties of networks representing arbitrary connectivity patterns. The number of patterns grows rapidly with the number of subpopulations, however, and the comprehensive description of networks of arbitrary size is a combinatorial challenge. Because small network motifs are the “building blocks” of large networks (Milo *et al*. 2002), the derivation of their features can be a step in predicting properties of complex connectivity networks. We thus characterize coalescent quantities under all possible motifs describing the spatial arrangements of up to four subpopulations. We first derive the expected coalescence times between pairs of lineages sampled in each of the subpopulations and pairs sampled from different subpopulations. For each subpopulation, we compute three population-genetic quantities: expected nucleotide diversity, expected *F*_*ST*_ values between pairs of subpopulations, and half-time to equilibrium after a perturbation. For each motif, we compute four network statistics—number of vertices, number of edges, mean degree, and density—correlating them with the population-genetic quantities. Finally, we investigate the nucleotide diversity lost after a connectivity loss or a subpopulation loss—a transition between motifs. We interpret the results in relation to problems in conservation genetics, considering two case studies, birds of genus Brachypteryx and Indian tigers. For both examples, we (i) consider genetic data in a network motif framework, and (ii) evaluate the potential impacts of connectivity change on population-genetic variation.

## Model

### Population connectivity

We consider *K* haploid or diploid subpopulations of equal size *N* individuals. We denote by *M*_*ij*_ the scaled backward migration rate, representing twice the number of lineages per generation from subpopulation *i* that originate from subpopulation *j*. Thus, *M*_*ij*_ = 2*Nm*_*ij*_ for haploids and 4N*m*_*ij*_ for diploids, where *m*_*ij*_ is the probability for a lineage of subpopulation *i* to originate from subpopulation *j* in the previous generation. The total scaled migration rate of subpopulation *i*, or twice the scaled number of lineages that originate elsewhere, is 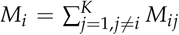. We further assume that the numbers of migrants from each non-isolated subpopulation are all equal to *M*, so that for two non-isolated subpopulations *i* and *j*, *M*_*i*_ = *M*_*j*_ = *M*. Time is a continuous variable *t*, scaled in units of the size of a single subpopulation (*N* for haploids, 2*N* for diploids). We focus on cases with 1 ≤ *K* ≤ 4, and we consider all possible connectivity patterns between subpopulations, where each pattern represents a distinct graph on at most four vertices (Figure 1).

### Coalescence

We consider the fate of two gene lineages drawn from a specific pair of subpopulations, either the same or different subpopulations. We denote the state of the two lineages by (*ij*), where *i* and *j* correspond to subpopulations. As the coalescence times between two lineages with initial states (*ij*) and (*ji*) are the same, we consider that (*ij*) refers to both (*ij*) and (*ji*), and we assume without loss of generality that *i* ≤ *j*. Consequently, the number of states for two lineages in *K* subpopulations is 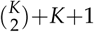: this quantity includes 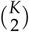 states where they are in different subpopulations, *K* where they are in the same subpopulation, and 1 state where they have coalesced.

Assuming that events cannot occur simultaneously, the coalescent process can be described by a continuous-time Markov chain (Kingman 1982; Wilkinson-Herbots 1998). The list of all possible states of the Markov chain in the case where *K* = 3 is represented in Figure 2.

**Figure 2.**
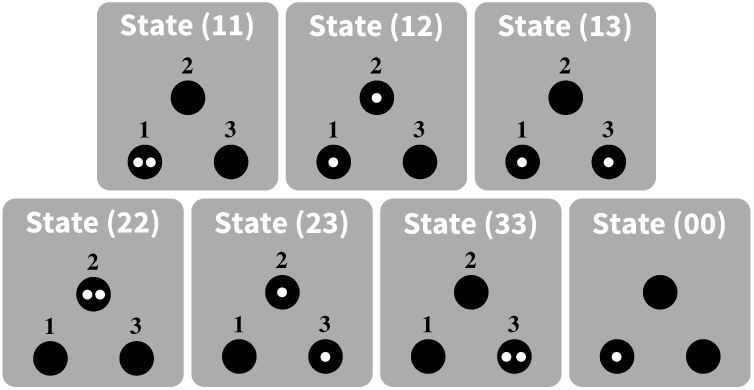
Schematic representation of all states for two lineage in a population divided into *K* = 3 distinguishable subpopulations. Lineages appear in white, and subpopulations appear in black. The two lineages can either be in different subpopulations (states (12), (13), and (23)), in the same subpopulation ((11), (22), and (33)), or they can already have coalesced ((00)).

The instantaneous rate matrix *Q* = (*q*_*ij, kℓ*_) for the Markov chain, where *q*_*ij, kℓ*_ is the instantaneous transition rate from state (*ij*) to state (*kℓ*), is defined by (Wilkinson-Herbots 1998):

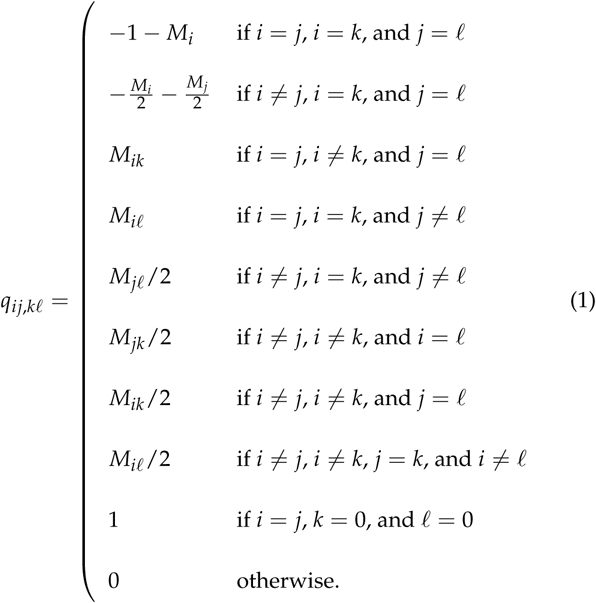

It can be seen that the list in eq. 1 covers all cases for (*i*, *j*, *k*, *ℓ*) by noting that by assumption, *i* ≤ *j* and *k* ≤ *ℓ*.

The transition probabilities between states after a time interval of length t are given by

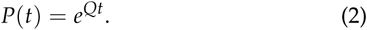

The element *p*_*ij, kℓ*_(*t*) of *P*(*t*) corresponds to the transition probability from state *ij* to state *kℓ* in time *t*.

This general model embeds known models. Setting *M*_*ij*_ = *M*/(*K* – 1) for all *i* and *j* ≠ *i* leads to the finite island model (Notohara 1990; Nei and Takahata 1993). Considering subpopulations along a circle and setting *M*_*ij*_ = *M*/2 for all adjacent subpopulations (*i* = *j* + 1, *i* = *j* – 1, or {*i*, *j*} = {1, *K*}) and *M*_*ij*_ = 0 for all non-adjacent subpopulations leads to the circular stepping-stone model (Strobeck 1987). Considering subpopulations along a finite line and setting *M*_*ij*_ = *M*/2 for 1 < *i* < *K*, *M*_12_ = *M*_*K*, *K*–1_ = *M*, and *M*_*ij*_ = 0 for all non-adjacent subpopulations leads to the linear stepping-stone model (Wilkinson-Herbots 1998).

## Results

### Expected coalescence time

The probability that coalescence has already occurred after time *t* for two lineages sampled respectively in subpopulations *i* and *j* corresponds to the transition probability during time t from initial state (*ij*) to state (00). This probability is given by element *P*_*ij*,00_ from matrix *P*(*t*) (eq. 2). Because *p*_*ij*,00_(*t*) is a cumulative probability, the associated density function is

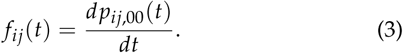

The expected coalescence time for two lineages sampled in sub-populations *i* and *j* is thus

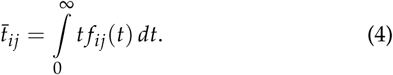

We derive in Appendix A the system of equations that can be solved to obtain the expected coalescence times in cases with one to four subpopulations. The expected coalescence times for motif 1—one isolated subpopulation—is simply 1. The expected coalescence times for the two-vertex motifs (motifs 2 and 3) appear in Table 1, for the three-vertex motifs (4 to 7) in Table 2, and for the four-vertex motifs (8 to 18) in Table 3.

**Table 1.** Exact mean coalescence times and *F*_*ST*_ values for 2-vertex motifs. *t*̄_*ij*_ represents the expected coalescence time for a pair of lineages, one sampled from subpopulation *i* and one sampled from subpopulation *j* (eq. 4). *F*_*ij*_ is the value of *F*_*ST*_ between subpopulations *i* and *j* (eq. 6).

| Motif | $\bar{t}_{11}$ | $\bar{t}_{22}$ | $\bar{t}_{12}$ | $F_{12}$ |
| --- | --- | --- | --- | --- |
| 2 | 1 | 1 | ∞ | 1 |
| 3 | 2 | 2 | $2(1 + \frac{1}{2M})$ | $\frac{1}{1+4M}$ |

**Table 2.**
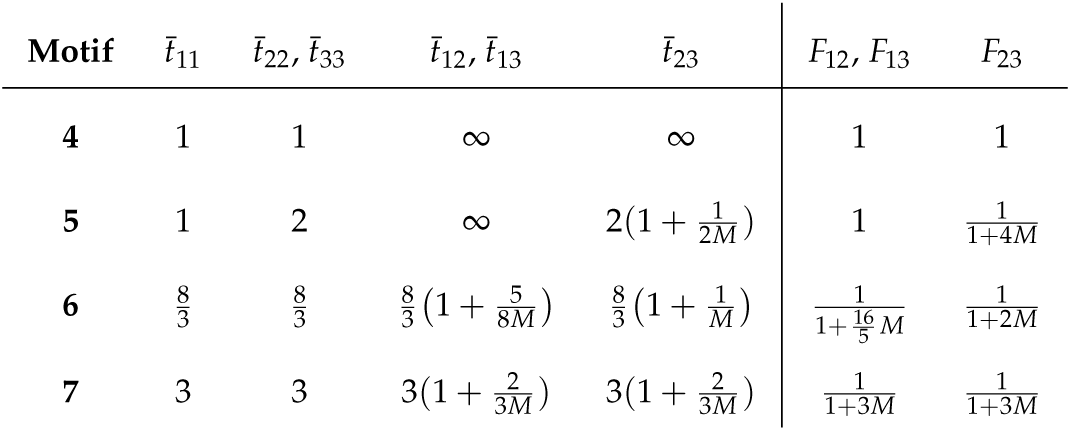
Exact mean coalescence times and *F*_*ST*_values for 3-vertex motifs. Owing to symmetries in migration motifs (Figure 1), *t*̄_22_ = *t*̄_33_ and *t*̄_12_ = *t*̄_13_, and thus, *F*_12_ = *F*_13_.

**Table 3.**
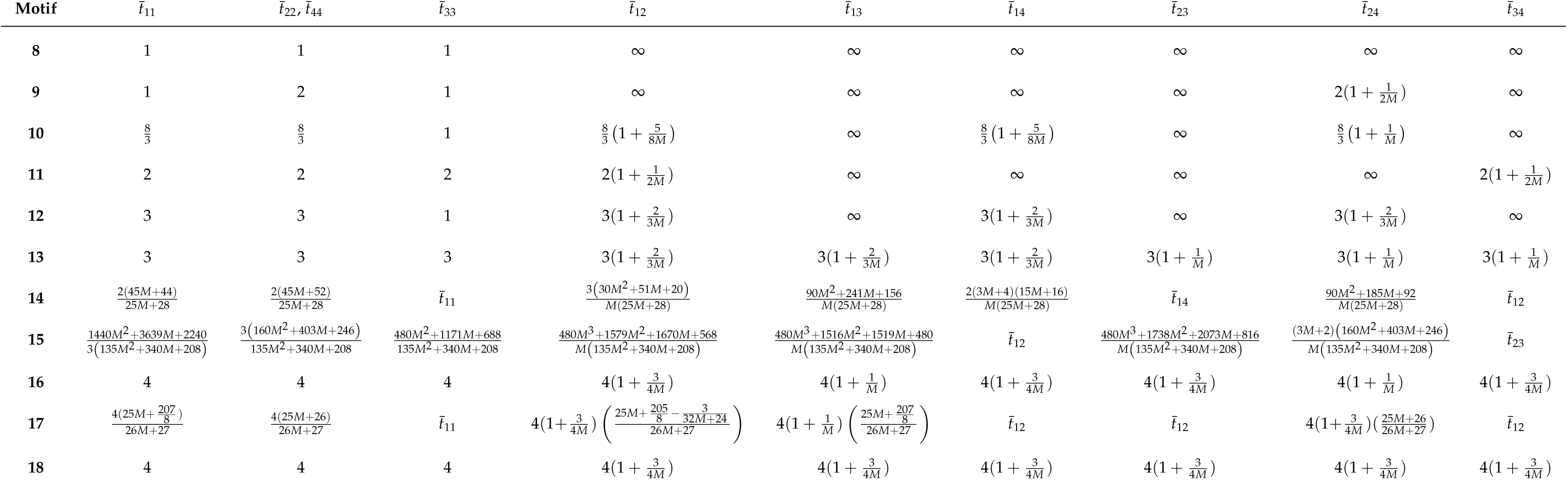
Exact mean coalescence times for 4-vertex motifs. Owing to symmetries in migration motifs (Figure 1), *t*̄_22_ = *t*̄_44_.

The set of all pairwise coalescence times of a motif is informative about another quantity of interest: the total coalescence time, that is, the coalescence time of two lineages randomly sampled in any two *K* subpopulations, possibly the same one. Indeed, the total coalescence time is simply 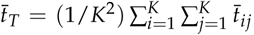, the mean coalescence time across all possible subpopulation pairs. The total coalescence times for all motifs presented in Figure 1 appear in Table S1.

### Expected within-subpopulation nucleotide diversity

We next calculate the expected within-subpopulation nucleotide diversity, that is, the expected number of differences between two nucleotide sequences sampled from the same subpopulation, assuming an infinitely-many-sites model (Kimura 1969) and a scaled mutation rate *θ* per site per generation. Here, *θ* represents twice the number of mutant lineages per generation in a subpopulation (2*Nμ* for haploids, 4*Nμ* for diploids, where *μ* is the unscaled per-site per-generation mutation rate). We take the mean across all subpopulations of the pairwise coalescence time within subpopulations:

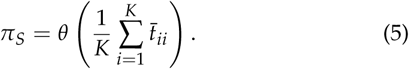

Note that *π*_*S*_ is also informative about total nucleotide diversity when *M* is large, because from Tables 1-3 and S1, the total coalescence time tends to the mean within-subpopulation coalescence time across all subpopulations as *M* → ∞.

We analytically computed the within-subpopulation nucleotide diversities for each motif by substituting the expected coalescence time from Tables 1-3 into eq. 5. Nucleotide diversity appears in Figure S1 as a function of network metrics.

### Genetic differentiation

For each motif, we compute expected values of *F*_*ST*_ between pairs of distinct subpopulations *i* and *j*, denoted by *F*_*j*_, from pairwise coalescence times. From Slatkin (1991),

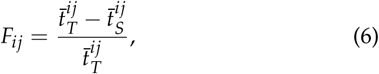

where 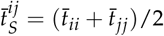 is the expected coalescence time of two lineages sampled in the same subpopulation, and 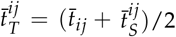 is the expected coalescence time of two lineages sampled in the total population. We compute eq. 6 using eq. 4.

For a *K*-vertex motif, *F*_*ST*_ has mean

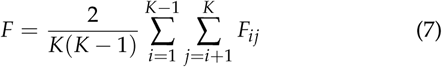

across subpopulation pairs. We analytically computed the expected *F*_*ST*_ from eq. 7 for each motif for sets of 3 and 4 subpopulations (Figure 1). The expected pairwise *F*_*ST*_ values for 2-, 3-, and 4-vertex motifs appear in Tables 1,2, and 4, respectively. *F*_*ST*_ appears in Figure S1 as a function of network metrics.

**Table 4.**
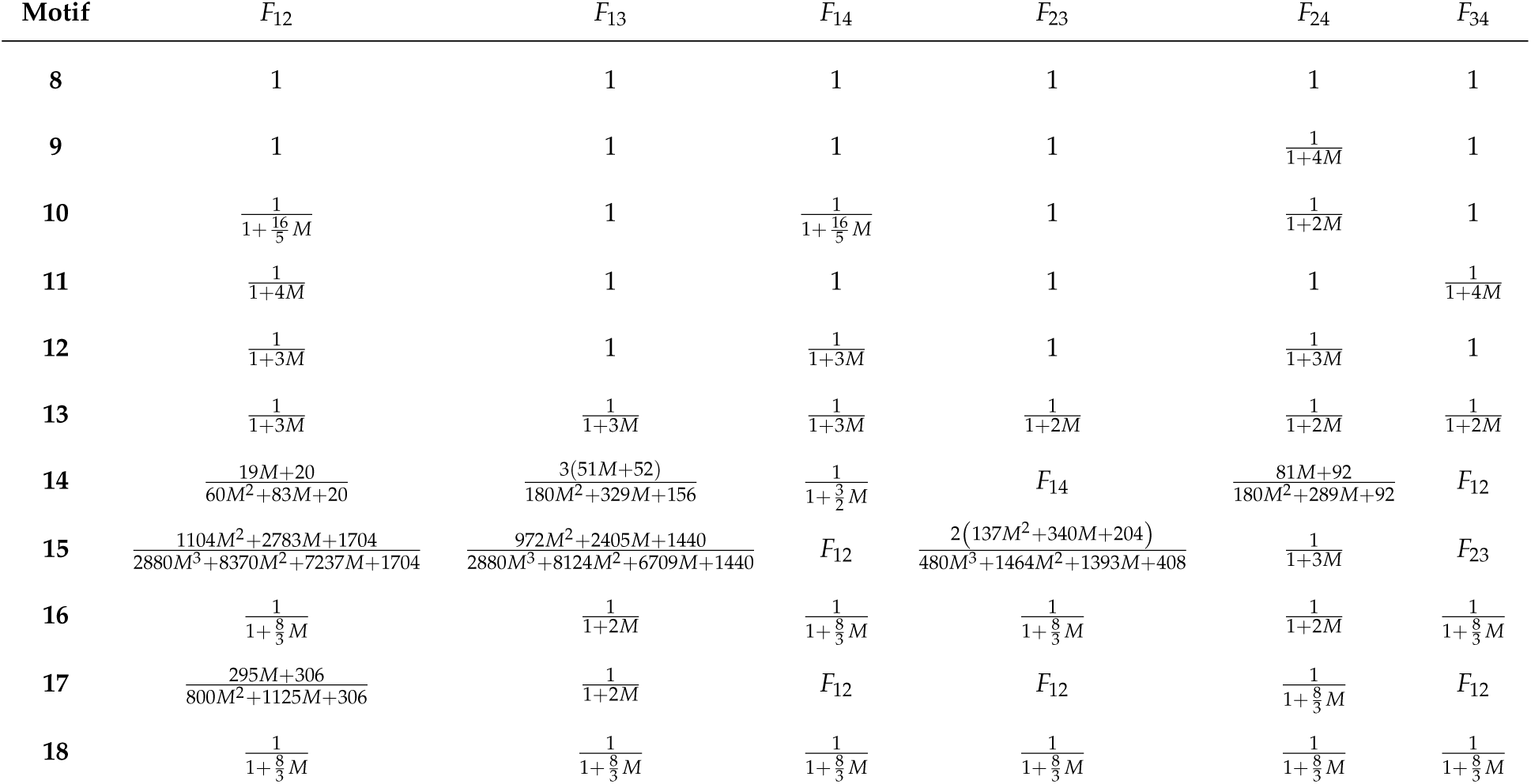
Exact *F*_*ST*_ values for 4-vertex motifs.

### Half-time to equilibrium diversity

The dynamics of *π*_*S*_ and *F*_*ST*_ are governed by the eigenvalues of matrix *Q* (eq. 1; e.g., Slatkin 1991). Considering an event that changed the population demography *τ* time units ago, *π*_*S*_ and *F*_*ST*_ will be at equilibrium in the sense that their values are stable through time if the probability that coalescence occurs at time *t* > *τ* is small, and thus, if *P*(*τ*) = *e*^Qτ^ ≈ [0,0,…, 0,1]^*T*^, where the last entry corresponds to the coalesced state.

Considering the eigendecomposition *Q* = *U*Λ*U*^-1^, where Λ is the diagonal matrix whose elements correspond to the eigenvalues of *Q* and *U* is the matrix whose columns are the eigenvectors of *Q*, *P*(*τ*) = *Ue*^Λ*τ*^*U*^‒1^. Thus, *P*(*τ*) ≈ [0,0,…, 0,1]^*T*^ when *e*^Λ*τ*^ ≈ [0,0,…, 0,1]^*T*^, which requires that *e*^*λ*_*i*_*τ*^ ≈ 0 for all eigenvalues *λ*_*i*_ except one, for which *e*^*λ*_*i*_*τ*^ ≈ 1. This condition holds if the largest eigenvalue of *Q* is 0 and the second-largest—denoted by *λ*—satisfies *e*^*λτ*^ ≈ 0. Because *Q* is an irreducible instantaneous rate matrix, its largest eigenvalue is 0 and all other eigenvalues are strictly negative (corollary 4.9 in Asmussen 2008).

We define the half-time to equilibrium *τ* as a function of *λ*, the second-largest eigenvalue of matrix *Q*, as

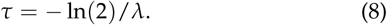

*τ* corresponds to the time at which *e*^*λτ*^ = 1/2. Thus, when *t* ≫ *τ*, *P*(*t*) ≈ [0,0,…, 0,1]^*T*^, and *π*_*S*_ and *F*_*ST*_ are approximately at equilibrium. The value of *τ* gives a sense of the time needed for *π*_*S*_ and *F*_*ST*_ to reach equilibrium values after a perturbation, such as after a loss of a connection or a subpopulation. This value depends on subpopulation connectivity patterns.

We computed the half-time to equilibrium from eq. 8 for each motif for sets of 1 to 4 subpopulations (Figure 1), numerically evaluating the second-largest eigenvalue of *Q*. Results appear in Figure S1 as a function of network metrics.

### Network motifs and patterns of genetic variation

To describe the influence of the properties of network motifs on our genetic variation measures, we computed the correlations between four network metrics and the mean within-subpopulation diversity *π*_*S*_, the mean *F*_*ST*_ across pairs of subpopulations, *F*, and the half-time to equilibrium diversity *τ*.

#### Network metrics

For a given motif, we denote by *V* and *E* its sets of vertices and edges, so that |*V*| and |*E*| correspond to the numbers of vertices and edges of the motif.

The first network metric we use is |*V*|, the motif size, or number of subpopulations *K*; here, |*V*| ranges from 1 to 4. The second metric is |*E*|, which corresponds to the number of pairs of subpopulations between which gene flow occurs; |*E*| ranges between 0 and 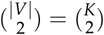. Our third metric is the mean vertex degree |*E*|/|*V*|, or the number of connections of an average subpopulation; it ranges from 0 to *K* – 1. The fourth network metric is the density 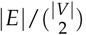, the number of edges divided by the maximum number of edges possible if the motif were a fully connected graph; it ranges from 0 to 1.

#### Correlations between network metrics and patterns of genetic variation

Correlations between network metrics and *π*_*S*_, *F*, and *τ* for motifs with up to four subpopulations appear in Figure 3. Diversity *π*_*S*_ is positively correlated with all four metrics, most strongly with the number of edges |*E*| (*ρ* = 0.96 for *M* = 10; Figure 3A) and the mean degree |*E*|/|*V*| (*ρ* = 0.96 for *M* = 0.1 and *M* = 1; Figure 3A). Indeed, the highest values of *π*_*S*_ occur for motifs 16, 17, and 18, which have the largest mean degree (2, 2.5, and 3, respectively), whereas the lowest values occur for motifs 1, 2, 4, and 8, which have mean degree 0.

**Figure 3.**
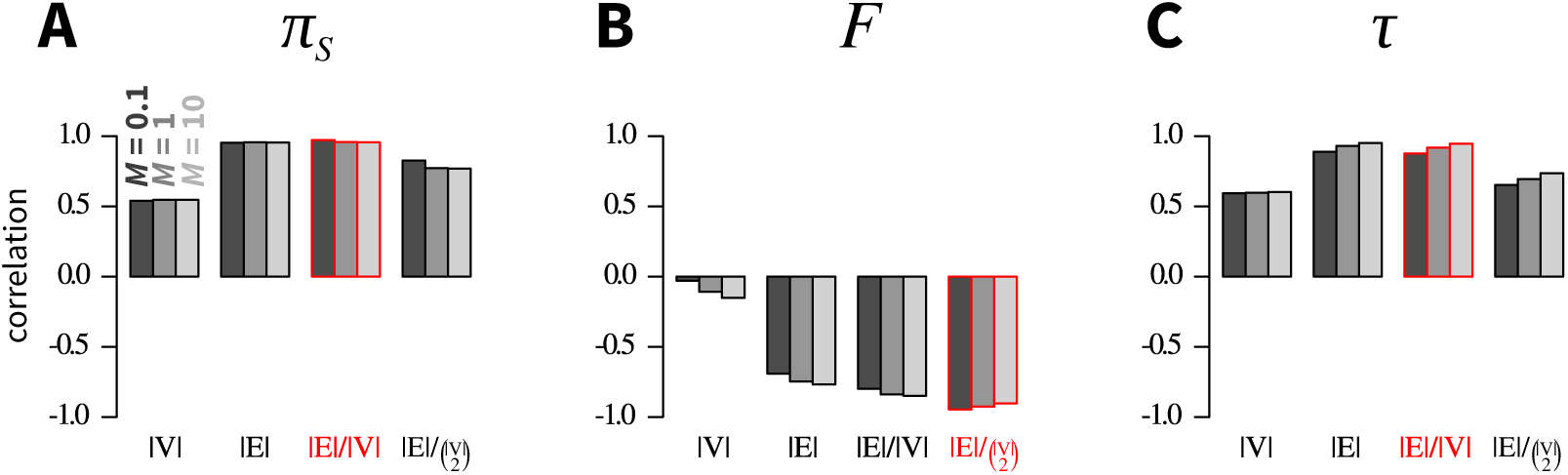
Pearson correlations between network metrics and genetic diversity measures. (A) *π*_*S*_, mean within-subpopulation nucleotide diversity (eq. 5). (B) *F*, mean pairwise FST across subpopulations (eq. 7). (C) *τ*, half-time to equilibrium diversity (eq. 8). Network metrics include number of vertices |*V*|, number of edges |*E*|, mean number of edges per vertex |*E*|/|*V*|, and density of edges 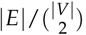. All network motifs in Figure 1 are considered. In each panel, the most strongly correlated metric appears in red.

*F* correlates negatively with the four metrics, especially the density 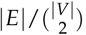 (*ρ* = –0.92 for *M* = 10; Figure 3B). Indeed, for large *M* (Figure S1H), the lowest *F* values occur for the densest motifs—3, 7, and 18—which have the maximal number of connections. The largest F values occur for the least dense motifs—2, 4, and 8—which have 0 edges.

Finally, *τ* is positively correlated with the four metrics, and most strongly with the mean degree |*E*|/|*V*| (*ρ* = 0.95 for *M* = 10; Figure 3C). For large *M* (Figure S1I), the largest *τ* values correspond to the motifs with largest mean degree (16,17, and 18), whereas the lowest *τ* values occur for the motifs with the lowest degree (1, 2, 4, and 8).

### Impact of a disturbance event

In this section, we focus on the impact of a disturbance event on mean genetic diversity *π*_*S*_. Of the three quantities we computed—*π*_*S*_, *F*, and *τ*—this quantity is perhaps the most central to conservation biology.

#### Enumerating outcomes of disturbance events

We enumerate all possible outcomes that could follow a disturbance event that removes a connection between two subpopulations or that removes a subpopulation. To do so, we compute a “graph of motifs,” where each vertex represents a motif, and we draw an edge between two motifs if they differ by a single subpopulation or a single connection. We orient edges of this graph from the motif with the larger number of subpopulations or connections toward the motif with the smaller number of subpopulations or connections. We give each edge a weight corresponding to the proportion of within-subpopulation diversity change associated with the transition from motif *i* to j, 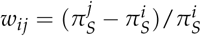, where 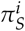 is the mean within-subpopulation diversity computed from eq. 5 applied to motif *i*. A negative weight indicates that the transition from motif *i* to motif *j* induces a loss of mean within-subpopulation diversity, whereas a positive weight indicates that the transition from motif *i* to motif *j* induces a gain of mean within-subpopulation diversity. In the case of a vertex loss, we consider that the lost subpopulation has diversity 0; for example, the transition from motif 3, where two subpopulations each have diversity 2 (Table 1), to motif 1, where a single subpopulation has diversity 1 and the “other” has diversity 0, leads to a change of *w*_20_ = [(1 + 0)/2 – (2 + 2)/2]/[(2 + 2)/2] = –0.75, that is, of 75% of the within-subpopulation diversity.

#### Edge losses and vertex losses

The graph of motifs appears in Figure 4A for edge loss and in Figure 4D for vertex loss. We focus on the case of *M* = 1.

**Figure 4.**
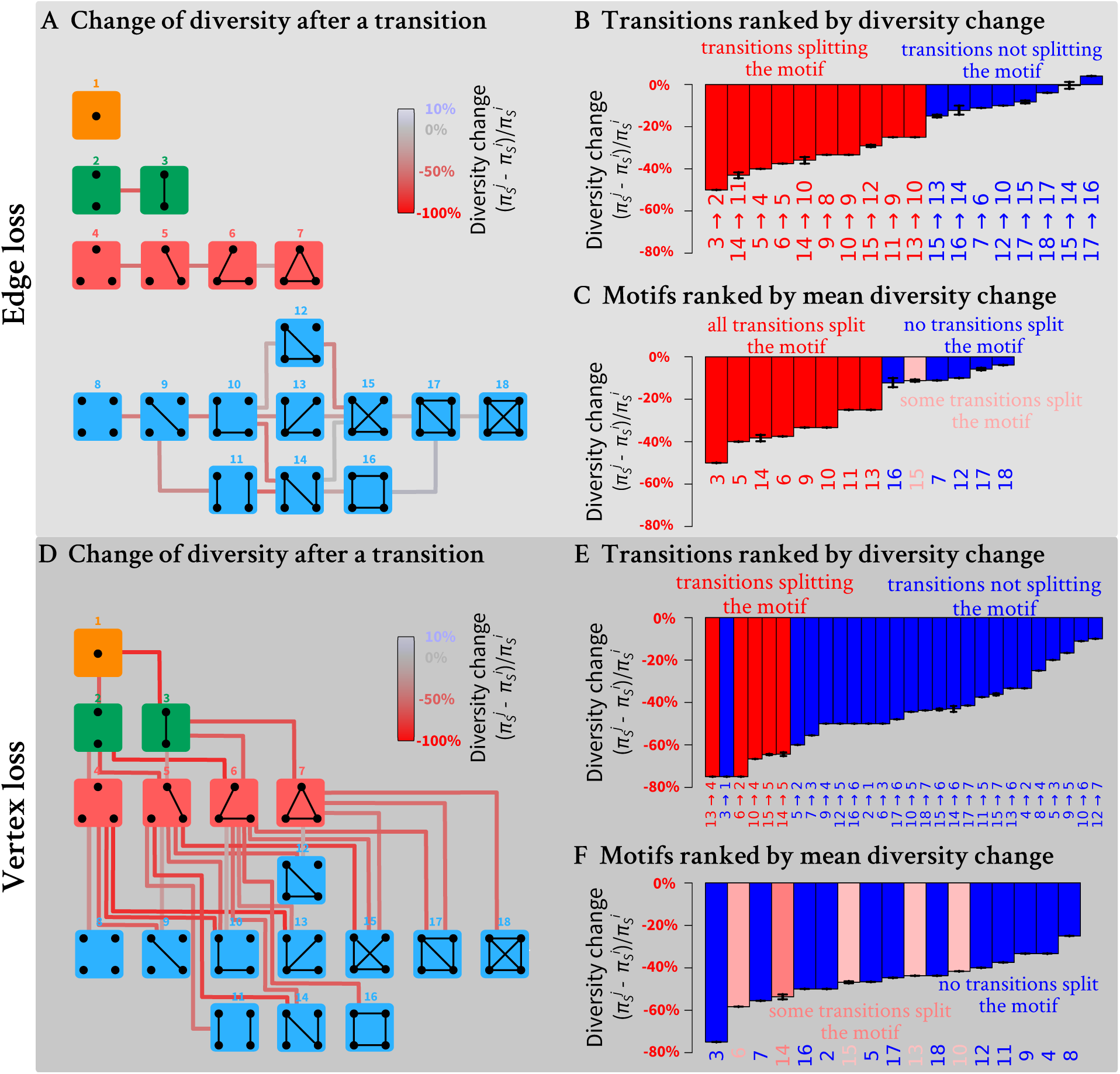
Change of within-subpopulation nucleotide diversity 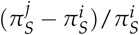 following a transition from motif *i* to motif *j*, for all possible transitions involving the loss of a single edge or a single vertex. (A) Schematic representation of all possible motif transitions involving an edge loss. Lines connecting motifs represent edge losses and are colored by changes of within-subpopulation nucleotide diversity: red for loss and blue for gain (see legend). (B) Motif transitions involving an edge loss ranked from the largest loss to the largest gain of within-subpopulation diversity. (C) Motifs ranked from largest to smallest mean diversity loss following edge loss. For each motif, the mean loss or gain is computed across all possible transitions to another motif. For example, for motif 5, three subpopulations can be lost; loss of the isolated subpopulation produces motif 3, generating a diversity loss of 20%, and loss of one of the two other subpopulations produces motif 2 and a diversity loss of 60%. Therefore, the mean diversity loss for motif 5 is (20%+60%+60%) /3 ≈ 46.7%. (D) Schematic representation of all possible motif transitions involving a vertex loss. Lines connecting motifs represent vertex losses and are colored by changes of within-subpopulation nucleotide diversity. (E) Motif transitions involving a vertex loss ranked from the largest to the smallest diversity within-subpopulation diversity loss. (F) Motifs ranked from largest to smallest mean diversity loss following vertex loss. In all panels, 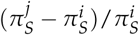 values assume *M* = 1; in (B), (C), (E), and (F), black horizontal bars represent minimum and maximum values of 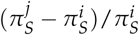 for *M* in (0, ∞). Values of 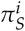 and 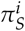 are computed from eq. 5 using coalescence times *t*̄_*ii*_ from Tables 1-3; minima and maxima of 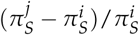 are obtained numerically. Pink shaded bars in (B), (C), (E), and (F) indicate that a fraction of edge losses or vertex losses split a motif: 1/4 for edge loss from motif 15 or vertex loss from motif 10,13, or 15,1/3 for vertex loss from motif 6, and 1/2 for vertex loss from motif

Loss of an edge can lead to diversity changes ranging from a loss of 50% to a gain of 4% (Figure 4B). Interestingly, the tran-sitions that lead to the greatest losses all split a motif into dis-connected sets of subpopulations (transitions in red, Figure 4B). The greatest diversity loss occurs with the transition from motif 3—which has a single connected pair of subpopulations—to motif 2—which has two isolated subpopulations. Surprisingly, one edge-loss transition increases the diversity for all migration rates in (0, ∞): the transition from motif 17 to motif 16. This transition increases the coalescence time for lineages sampled from different subpopulations without isolating any subpopulations.

The impact on diversity of the loss of a vertex ranges from a loss of 75% to a loss of 10% (Figure 4E). Similarly to the edge loss case, the vertex losses that lead to the greatest losses generally correspond to a split of the motif into disconnected sets of subpopulations (transitions in red, Figure 4E). For instance, the greatest diversity loss is associated with the transition from motif 13—which has a single set of four connected subpopulations—to motif 4—which has three isolated subpopulations.

#### Fragile and robust motifs

We can also identify the most “fragile” motifs: the motifs for which disturbance leads to the greatest diversity loss. For each motif, we compute the diversity changes associated with all |*E*| edge or |*V*| vertex losses, reporting the mean across the edge or vertex set. Motifs ranked by robustness to an edge loss appear in Figure 4C. The most fragile motifs are those split into disconnected components by an edge loss, whereas the most “robust” motifs are those that are not split.

Motifs ranked by robustness to a vertex loss appear in Figure 4F. We can see that the most fragile motifs are motifs 3, 6, and 14 (linear stepping-stone models) and motifs 7 and 16 (circular stepping-stone models). The linear stepping-stone motifs are easily split by a vertex loss, producing a disconnection that is expected to reduce diversity. The circular stepping-stone models, however, are not easily split by a vertex loss. Their fragility stems from their high diversity, among the highest of all models, on par with island models (Tables 1, 2, and 3). Any motif transition is thus likely to substantially reduce diversity.

## Examples

We use the results from our network-based model to reinterpret spatial genetic structure in two animal examples. Using published spatial and genetic information for each example, we propose a network motif that might represent the structure of the population. We then ask what types of transitions could result in increased or decreased population structure and variation in the context of the conservation biology of the species examined.

### Indian sky island birds of genus Brachypteryx

First, we consider two species of genus *Brachypteryx*, birds endemic to the Western Ghats sky islands of India: the white-bellied and the rufous-bellied shortwings *Brachypteryx albiventris* and *B. major*. Robin *et al*. (2015) reported microsatellite data from multiple geographically separated subpopulations, sampling 218 individuals at 14 microsatellite loci. These subpopulations have experienced changes in geographic range and gene flow on both evolutionary and anthropogenic time scales owing to Pleistocene climate change that could have shifted the locations of suitable habitat and recent deforestation. Such changes can influence numbers of populations and gene flow between them, and can be interpreted using our network model.

Robin *et al*. (2015) stated that genetic differentiation in the pair of species was not quite consistent with a simple island-migration model, so that our network approach might provide additional insight. Indeed, consistent with geographic barriers, Robin *et al*. (2015) observed genetic differentiation between the two species, as well as two subgroups within each species. The data generally fit motif 11 (Figure 5A), containing two relatively isolated sets of subpopulations, each with two subpopulations that exchange migrants. However, *F*_*ST*_ values between the two species sets (Table S3 of Robin *et al*. 2015) were lower than the high values expected under motif 11 (Table 4), potentially as a result of a short time scale of fragmentation.

**Figure 5.**
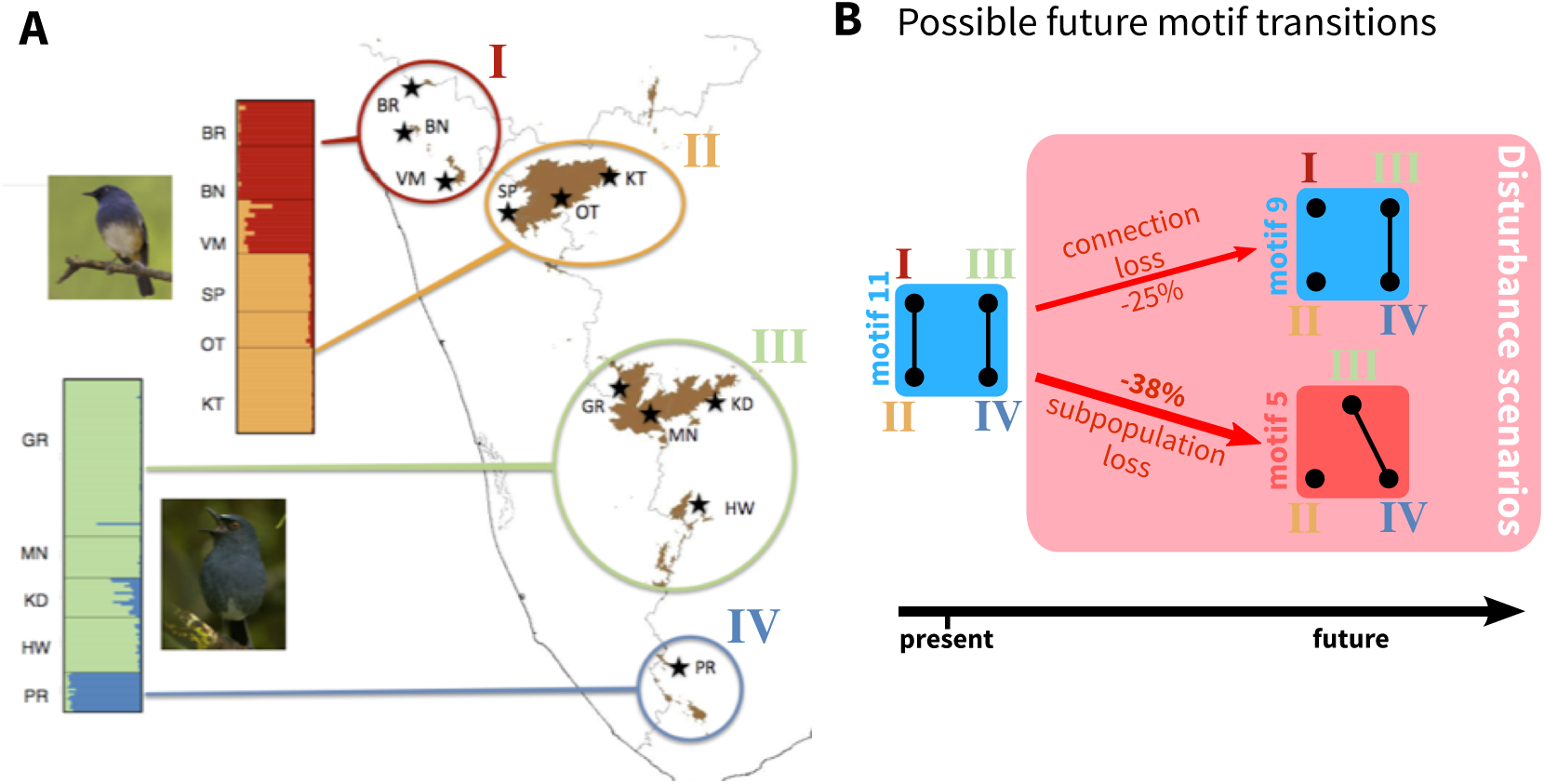
Application of the network theory framework to the Indian sky island birds *Brachypteryx albiventris* and *B. major*. (A) Map of the distribution of *B. albiventris* and *B. major* in the Indian sky islands of the Western Ghats with sampling locations, and STRUCTURE plot. The map and the STRUCTURE plot are adapted from Robin *et al*. (2015). Colors and roman numerals represent the four genetic clusters. Two-letter codes indicate the sampling locations. Sampling locations for *B. major* include BR: Brahmagiri; BN: Banasura; VM: Vellarimala; SP: Sispara; OT: Ooty; KT: Kothagiri. Sampling locations for *B. albiventris* include GR: Grasshills; MN: Munnar; KD: Kodaik-anal; HW: High Wavys; and PR: Peppara. (B) Possible future motif transitions, based on the transitions from motif 11, which is taken to represent the current state of the population. Numbers on arrows represent predicted losses of mean nucleotide diversity across subpopulations (Figure 4).

Under the network model, supposing that the current motif is 11, we can investigate the future impact of the loss of an edge or vertex, representing events possible for an endangered species (Figure 5B). The transition from motif 11 to motif 9 is seen as a loss of an edge, corresponding to a loss of migration between one of the pairs of subpopulations. This event decreases within-subpopulation nucleotide diversity (-25%; Figure 4C), and leads to increasing *F*_*ST*_ genetic differentiation between subpopulations, particularly within each species. The loss of a subpopulation, transitioning from motif 11 to motif 5, similarly leads to a loss of within-subpopulation nucleotide diversity (-38%; Figure 4F).

Note that the losses reported are expected losses in the long term. The half-time to equilibrium τ values for motifs 5 and

9 appear in Figure S1C, F, I. Interestingly, they are equal, and correspond to 7.69 for *M* = 0.1, 1.81 for *M* = 1, and 1.42 for *M* = 10, in units of 2*N* generations. Thus, depending on the migration rate, the future decrease of genetic diversity substantially changes. The identical *τ* values for the two motifs result from the fact that *τ* is determined by the motif component with the lowest half time to equilibrium, and the two motifs have similar components—a pair of connected subpopulations and either one or two isolated subpopulations.

Comparing the edge loss and vertex loss scenarios, a vertex loss transition from motif 11 to motif 5 has a greater negative effect on nucleotide diversity, because it has the largest longterm effects and the equilibrium is reached as quickly as in the edge loss transition. In this case, focusing on preserving subpopulations rather than gene flow is predicted to avoid the most detrimental loss of genetic diversity for the subpopulations.

### Indian tigers

Next, we consider genetic variation for tigers in India, representing 60% of the global wild tiger population (Mondol *et al*. 2009). Natesh *et al*. (in press) considered the genetic diversity and structure across the Indian subcontinent of tigers, a species that now occupies 7% of its historical range. India’s ~2,500 tigers are distributed across many small groups, with a median size of 19 across recognized groups. Understanding population structure and connectivity is important to tiger conservation.

Using 10,184 SNPs, Natesh *et al*. (in press) identified a north-western subpopulation (dark blue cluster *i* in Figure 6A), a north/northeastern subpopulation (green cluster II), a central subpopulation (orange cluster III), and a southern subpopulation (purple cluster IV). They reported evidence of gene flow between subpopulations III and IV and between subpopulations II and III. The exact relationship between subpopulations, however was unclear. From the pairwise *F*_*ST*_ values reported in Table 2 of Natesh *et al*. (in press), levels of divergence between subpopulation I and all other subpopulations were high, suggesting isolation with limited gene flow. fastSTRUCTURE analyses performed by Natesh *et al*. (in press) suggested connectivity between subpopulations II and III (Figure 6A). Owing to the large *F*_*ST*_ between the northeastern subpopulation (II) and the southern subpopulation (IV) and between the central subpopulation (III) and the southern subpopulation (IV), we suggest that the motif most clearly fitting the current population structure is motif 9.

**Figure 6.**
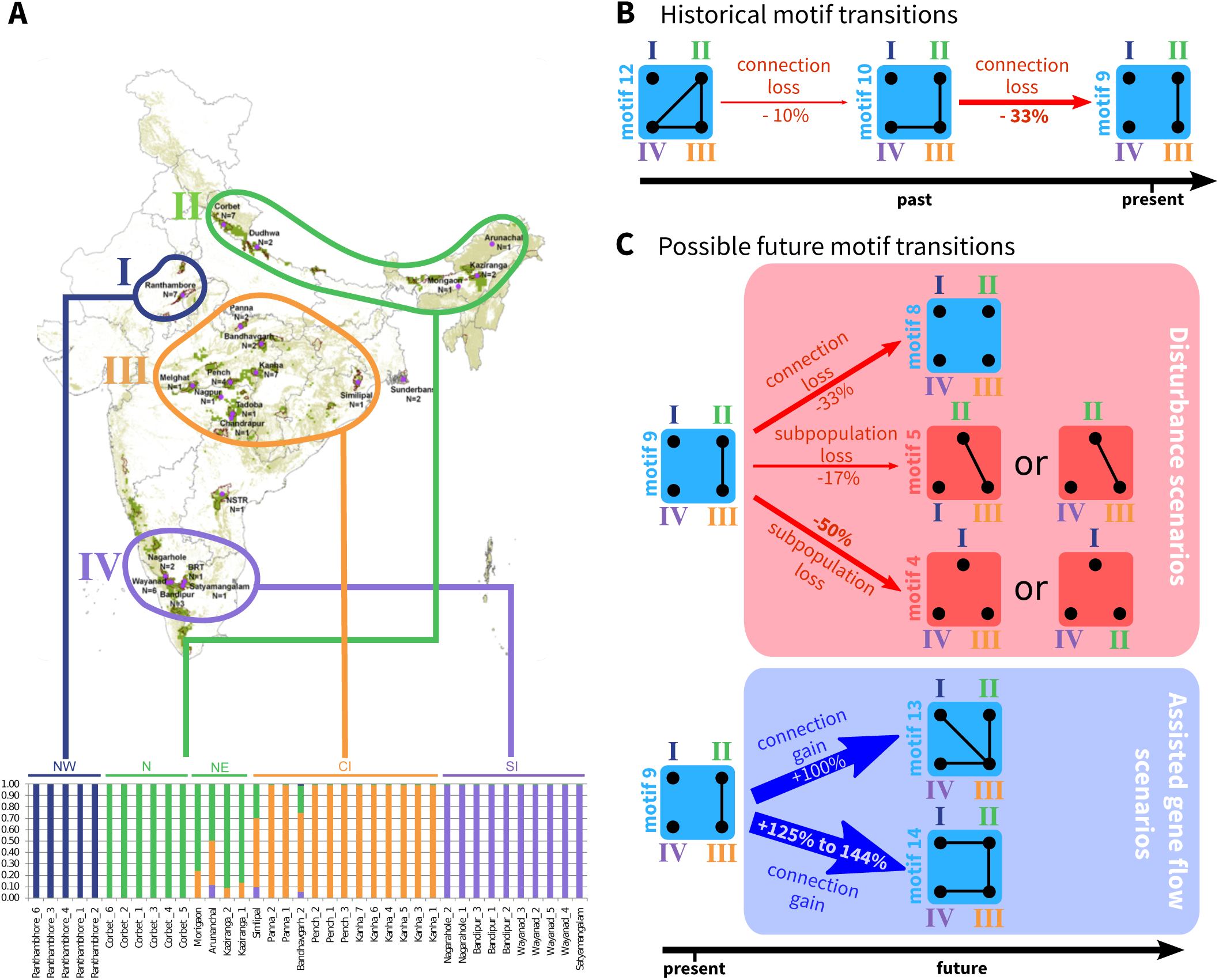
Application of the network theory framework to Indian tigers. (A) Map of the distribution of tigers with sampling locations, and fastSTRUCTURE plot. The figure is adapted from Natesh *et al*. (in press). Note that sample sizes for the fastSTRUCTURE plot include only subsets of individuals from Natesh *et al*. (in press). (B) Hypothetical sequence of past motif transitions based on Natesh *et al*. (in press) and Figure 4. (C) Possible future motif transitions, based on the transitions from motif 9 (Figure 4). For (B) and (C), motif 9 is taken to represent the current state of the population; percentages correspond to the proportion of within-subpopulation diversity change following each motif transition (from Figure 4).

Because of the smaller pairwise *F*_*ST*_ values between subpopulations II and IV and between subpopulations III and IV than between subpopulations I and IV, we suggest that a recent change in network structure occurred from motif 12 to motif 9 (Figure 6B), involving recent loss of connectivity between subpopulation IV and the other subpopulations, and leading to a loss of 40% of the within-subpopulation diversity (-10% from the transition from motif 12 to motif 10, and then −33% from the transition from motif 10 to motif 9). That connectivity loss might have occurred recently is supported by previous genetic and historical data: an earlier study with 10 microsatellite markers suggests an older transition between motif 15 and 12, with connectivity loss between subpopulations I and II (Mondol *et al*. 2013).

Ongoing perturbations to the network are likely, owing to increasing human pressures and land-use changes that reduce population sizes and increase fragmentation (Figure 6C). The transition from motif 9 to motif 8, involving the loss of an edge, would decrease within-subpopulation nucleotide diversity by 33% (Figures 4C and 6C). The loss of a subpopulation, however, leads to qualitative differences in the genetic structure depending on the subpopulation lost. If the more isolated northwestern subpopulation *i* or the southern subpopulation IV is lost, then the resulting network is similar to motif 5, with a moderate decrease in within-subpopulation nucleotide diversity (-17%) and a decrease in differentiation overall because an isolated subpopulation is lost (Figures 4C and 6C). By contrast, if one of the connected subpopulations, the central subpopulation III or the northern/northeastern subpopulation II, is lost, then a decrease in diversity is expected (-50%; Figures 4C and 6C).

To maintain or restore some of the recently lost genetic diversity of Indian tigers, Kelly and Phillips (2016) suggested reconnecting isolated subpopulations by assisted migration (Figure 6C). Two such reconnection scenarios can be imagined. The first scenario, which corresponds to restoring lost migration routes (Figure 4B), reconnects the central subpopulation III with all other subpopulations, producing a transition from motif 9 to motif 13. It would lead to an increase of within-subpopulation diversity of 100%. Alternatively, a second scenario in which subpopulations are reconnected along a line, forming a linear stepping-stone, is possible, corresponding to a transition from motif 9 to motif 14. This scenario might seem less intuitive, as it does not correspond to any previous population structure. Interestingly, it leads to a greater increase of diversity (+125% to +144%, depending on the amount of gene flow; Table S2).

Note that the losses and gains of diversity reported are expected losses and gains in the long term. The *τ* values for motifs 4, 5, 8, 13, and 14—the motifs that are possible as the result of the transitions in Figure 6C—appear in Figures S1C, S1F, and S1I. *τ* has the same value of 0.69 for both motifs 4 and 8, because they have only isolated subpopulations. *τ* for motif 5 is 7.69 for *M* = 0.1,1.81 for *M* = 1, and 1.42 for *M* = 10, in units of 2*N* generations. Thus, in addition to being the transition leading to the greatest diversity loss, the transition from motif 9 to motif 4 is also the one that affects diversity the fastest. *τ* for motif 13 is 17.76 for *M* = 0.1, 3.34 for *M* = 1, and 2.19 for *M* = 10, in units of 2N generations. *τ* for motif 14 is 29.32 for *M* = 0.1, 4.72 for *M* = 1, and 2.68 for *M* = 10, in units of 2*N* generations; the time required to restore diversity exceeds the time it takes to lose it. Among assisted gene flow scenarios, the transition to motif 14 that leads to the larger amount of diversity in the long term among the pair of scenarios considered is the one with the slower change of diversity. This result suggests a trade-off between the magnitude and speed of the transition to long-term effects on diversity.

## Discussion

We have presented a novel framework that combines network theory and population genetics to study the impact of population structure on patterns of genetic variation under diverse assumptions about population connectivity. Treating a structured population as a network containing vertices that represent subpopulations and edges that represent gene flow, considering all possible population network motifs for sets of one to four subpopulations, we have determined motif features that correlate with patterns of genetic variation. Among four motif statistics, we found that the mean node degree is the most strongly correlated with within-subpopulation diversity, and that motif density is the most strongly correlated with genetic differentiation between subpopulations.

Our framework makes it possible to predict the impact on genetic diversity of disturbances such as loss of a subpopulation or a connection between subpopulations. The effect of the loss of a vertex or edge depends on the context of the disturbance in the population network. Whereas some disturbances that split the network, including edge losses in transitions from motif 3 to 2 and from 14 to 11 and the vertex loss in the transition from motif 13 to 4, substantially reduce genetic diversity, others such as the transition from motif 17 to 16 instead increase mean diversity across subpopulations (Figure 4).

### Theoretical advances

Our results extend classical coalescent theory results concerning migration models. Among the 18 network motifs we studied, 11 correspond to migration models that differ from the standard models.

As has been seen previously (Slatkin 1987; Strobeck 1987; Wilkinson-Herbots 2003), for motifs all of whose subpopulations are exchangeable and none of whose subpopulations are isolated, we find that the within-subpopulation pairwise coalescence times are independent of the migration rate (Table S2). Interestingly, we found that this result on migration rate independence also holds for motifs with disconnected components (motifs 2, 4, 5, 8-12), even though disconnection leads to violation of the assumption of migration matrix irreducibility used in Slatkin (1987). This result can be explained by the fact that such motifs all involve juxtaposition of smaller motifs, each of which has exchangeable subpopulations, none of which are isolated. Consequently, even though motifs 2,4, 5, and 8-12 do not satisfy the assumptions used in Slatkin (1987), that each component of the motif satisfies them suffices to ensure the result on migration-rate independence.

Motifs 14,15, and 17, for example, do not have exchangeable vertices, nor can these motifs be decomposed into disconnected components that each have exchangeable vertices. Their within-subpopulation coalescence times do depend on the migration rate (Table S2). Nevertheless, within-subpopulation coalescence times of all motifs vary relatively little with the migration rate: the difference between the maximum and minimum values is less than 15% of the minimum (Table S2). We find that migration rates have only a small effect on within-subpopulation diversity in many spatial configurations.

Our results also extend classical theoretical results about genetic differentiation. Under the island model, *F*_*ST*_ follows a formula 1/(1 + *αM*), where the constant *α* > 0 determines the relative impact of drift and migration (Wright 1951; Nei and Takahata 1993), and it approximately follows 1/(1 + *αM*) under the stepping-stone model (Cox and Durrett 2002). For a fixed number of subpopulations *K*, among networks with all nodes connected, *α* is smallest under the island model and largest under the stepping-stone model (Cox and Durrett 2002).

We exhibit additional models under which pairwise *F*_*ST*_ follows 1/(1 + *αM*), where α is intermediate between that expected under the island and stepping-stone models (Table S3). All models with exchangeable vertices or decomposable into components each with exchangeable vertices (motifs 2-5, 7-9, 11, 12, 16, 18) have *F*_*ST*_ values that follow this formula. Interestingly, motif 13, which does not have exchangeable vertices and is not decomposable in this manner, also has an *F*_*ST*_ that follows such a formula, with *α* = 2 or 3. Its *α* values lie near those of the island-migration motif 18, with *α* = 8/3, and the linear stepping-stone motif 14, with α ranging from 1.129 to 3.2 across population pairs and across migration rates, and are also similar to that of the circular stepping-stone motif 16 (*α* = 2 or 8/3), although motif 16 has more connections.

Motifs 15 and 17 also have non-exchangeable vertices and are not decomposable, and they have *F*_*ST*_ values whose expressions involve rational functions of *M* (Table 4). We show in Table S3, however, that their *F*_*ST*_ values approximately follow an expression of the form 1/(1 + *αM*), with α ranging from 1.747 to 2.989; in addition, their α values are close to that of motif 16, with *α* = 2 or 8/3. Although motifs 4, 5, and 8-12 have at least two disconnected components and thus their global *F*_*ST*_ is equal to 1 irrespective of the value of *M*, their pairwise *F*_*ST*_ values for connected subpopulations do follow 1/(1 + *αM*), with α ranging from 2 to 4. Overall, our results highlight that the classical formula *F* = 1/(1 + *αM*) is a helpful approximation for all motifs with up to four subpopulations.

### Data applications

Our results provide a framework for interpreting empirical patterns of genetic diversity and differentiation, and for predicting future patterns. We have illustrated how they provide insight into two systems of conservation interest, Indian sky island birds of genus *Brachypteryx* and Indian tigers. After suggesting the most appropriate motif for each species and the sequence of transitions that might have led to the current motif, we enumerated future disturbances and highlighted the ones that would have the strongest long-term impact on genetic diversity within subpopulations. For tigers, we enumerated possible assisted gene flow scenarios and highlighted scenarios leading to the greatest long-term genetic diversity increase.

Many studies have focused on deducing population networks from genetic data in population genetics; most such applications have focused on clustering using community detection algorithms, without using a population-genetic model (Dyer and Nason 2004; Dyer 2015; Garroway *et al*. 2008; Rozenfeld *et al*. 2008; Ball *et al*. 2010; Munwes *et al*. 2010; Greenbaum *et al*. 2016). While these statistical approaches are appealing for making sense of complex datasets, the mechanistic models we consider are useful for providing predictions about genetic diversity patterns. We have demonstrated how simple network motifs can be deduced from cluster analyses and pairwise *F*_*ST*_ values and can then be used to predict the impact of future disturbances.

The network theory framework is promising for the analysis of natural populations whose spatial arrangements do not follow classical migration models. For example, river systems involve subpopulations arranged along a stream, leading to a motif with a linear arrangement such as 3, 6, and 14, or in different streams, leading to a star-shaped motif such as 13 (Morrissey and de Kerckhove 2009). Geographic barriers owing to mountains, valleys, and human occupation can isolate one (motifs 2, 5, 11 and 12) or several subpopulations (motifs 4, 8, and 9). Moreover, many landscapes present a specific zone with high resistance, for example owing to low habitat quality, leading to partial isolation of a subpopulation from a strongly connected set of subpopulations (motif 15).

Our exhaustive enumeration of motifs ensures that we can confront empirical data with expected patterns of genetic variation under any spatial arrangement. This enumeration can improve our ability to interpret genetic data, especially for threatened species, which typically present high fragmentation and are likely to undergo future disturbances resulting from further human-induced habitat loss or from conservation efforts such as assisted gene flow. The framework is also promising for conservation planning, because it suggests which connections or subpopulations are more important in contributing to genetic variation. Historical human impacts, ongoing urbanization, and habitat fragmentation are leading to species range collapse and population decline (e.g. carnivores; Ripple *et al*. 2014). Some species, such as the sky island birds of genus Brachypteryx, are specialized to habitats that are naturally patchy and isolated (Robin *et al*. 2015). Understanding the consequences of such patchiness from a network perspective can provide insights on mitigation for ongoing habitat fragmentation.

In species such as the Indian tiger, conservation might require management strategies that include assisted migration (Kelly and Phillips 2016). In such contexts, strategies can be designed for maximizing genetic variation, by giving the existing set of subpopulations the most favorable connections possible. For designing such strategies, our approach provides an alternative to spatially explicit landscape-genetic models focused on effects in physical space, enabling assessment of the potential genetic consequences of alternative network motifs.

### Extensions

Several assumptions of our model could be relaxed to make it more closely match natural systems. We only considered homogeneous subpopulations, with equal sizes and similar migration rates in all non-isolated subpopulations, and equilibrium genetic variation. Heterogeneous sizes are common in environments with varying habitat quality (Dias 1996), and migration rate differences are common in species that disperse passively, such as by currents or wind (Vuilleumier and Possingham 2006). Permitting heterogeneity would increase the number of motifs possible for fixed numbers of subpopulations, potentially introducing source-sink dynamics (Dias 1996). These dynamics are expected to influence robustness to loss of a connection or subpopulation: we expect nucleotide diversity to be robust to loss of a connection to a sink subpopulation or loss of a sink subpopulation itself, because such subpopulations might be small with relatively low nucleotide diversity. Conversely, we expect nucleotide diversity to be less robust to the loss of a source subpopulation, as these subpopulations are typically larger and more diverse.

Non-equilibrium genetic diversity is common in species that face frequent environmental changes, and it can result in transient levels of genetic variation that strongly differ from the equilibrium and that persist for many generations (Alcala *et al*. 2013; Alcala and Vuilleumier 2014). The expected diversities in Tables 1-4 correspond to long-term expectations, and give a sense of the *potential* of a given spatial configuration to permit large levels of genetic diversity. To assess the impact of a perturbation, long-term expectations must be contrasted with the time to reach them. We thus advocate computation of the half-time to equilibrium *τ*, which gives a sense of the time needed for nucleotide diversity and *F*_*ST*_ to approach their equilibrium values. Interestingly, we find that *τ* is strongly correlated with the mean vertex degree; it would be worthwhile to assess the potential of |*V*| as a predictor of *τ* in larger networks.

### Conclusion

This work is a step toward developing a general theory that links network topology and patterns of genetic variation. Small motifs are the building blocks of complex networks (Milo *et al*. 2002); thus, for large networks, counting the number of appearances of each 3‐ or 4-vertex motif can give an initial idea of the fine-scale structure of the population. The results we have derived make it possible to link this fine-scale structure to local patterns of variation. For example, if we find many instances of motifs 17 or 18, we might conclude that the network is dense and thus has large diversity, low *F*_*ST*_, long time to equilibrium after a perturbation and a high robustness to perturbation. On the other hand, if we find many instances of lower-density motifs 9,10, and 14, we might reach the opposite conclusions.

The detection of motifs that are overrepresented in certain types of network (e.g. ecological, neural, protein-interaction) has been used to identify network classes that share common properties despite describing different data types (Milo *et al*. 2002; Alon 2007). Further work could consider motifs that are overrepresented in population networks, to assess whether population networks have a shared “motif signature” or if certain networks are more common in certain habitats (e.g. marine, river, terrestrial). Such an approach could help identify similarities between population networks and other types of biological networks. Our results can potentially be extended to larger networks, and it could be assessed how global patterns in genetic diversity and *F*_*ST*_ can be predicted from information on the occurrence of small motifs. Such an extension will become increasingly valuable as more empirical studies sample genomic datasets from broad geographical scales with fine-grained sampling resolution.

## Acknowledgments

We thank G. Greenbaum for comments. Part of this work was completed when UR was a visitor at the Stanford Center for Computational, Evolutionary, and Human Genomics (CEHG).

We acknowledge support from NSF grant DBI-1458059, a CEHG postdoctoral fellowship, and Swiss National Science Foundation Early Postdoc.Mobility fellowship P2LAP3_161869.

## Appendix A. Deriving expected coalescence times

Expected coalescence times can be obtained by first-step analysis. The expected coalescence time of all states (*ij*) (eq. 4), where *i* ranges in [1,2,…, *K* – 1] and *j* ranges in [*i*, *i* + 1,…, *K*], can be decomposed into a sum of expected coalescence times (Notohara 1990; Wakeley 1998)

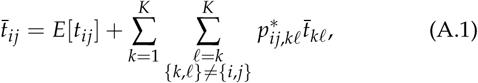

where *E*[*t*_*ij*_] is the expected time before a change of state, and

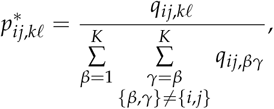

where *q* terms are taken from eq. 1. Eq. A.1 describes a system of 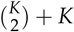 equations. In the next sections, we describe this system of equations in the cases of *K* = 1 to *K* = 4.

### 1-vertex motif

The case of one subpopulation has a single possible initial state, which is given by classical coalescent results (Kingman 1982):

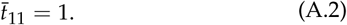

This quantity directly gives the expected pairwise coalescence time of motif 1.

### 2-vertex motifs

In the case of two subpopulations, *M*_12_ = *M*_21_ = *M*_1_ = *M*_2_ = *M*. Eq. A.1 then simplifies to

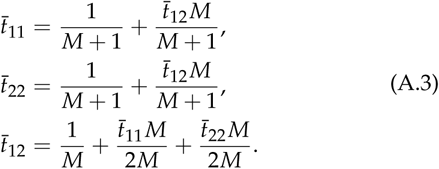

This system and its solution were derived in Nath and Griffiths (1993). Setting *M* = 0 and solving the system for *t*̄_11_, *t*̄_22_, and *t*̄_12_ gives the expected pairwise coalescence times of motif 2 (Table 1). Considering *M* > 0 and solving the system gives the expected pairwise coalescence times of motif 3 (Table 1).

### 3-vertex motifs

In the case of three subpopulations (Fig. A1), eq. A.1 becomes

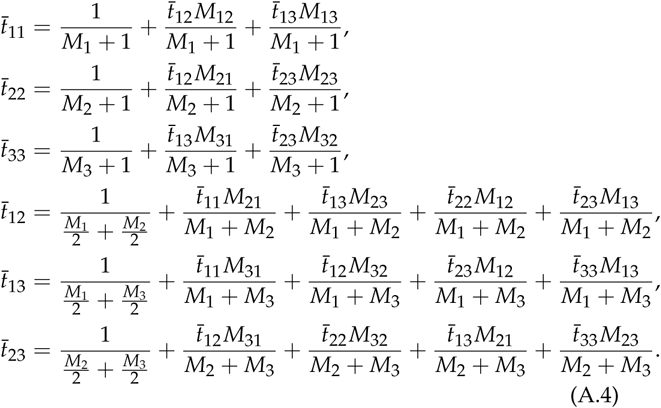

We set the values *M*_*ij*_ to reflect the network motifs of Figure 1, solve the linear system of equation, and report the corresponding expected times in Table 2. For example, for motif 5, *M*_1_ = 0, *M*_12_ = *M*_21_ = *M*_13_ = *M*_31_ = 0 and *M*_23_ = *M*_32_ = *M*_2_ = *M*_3_ = *M* in all equations, and we obtain the system of equations

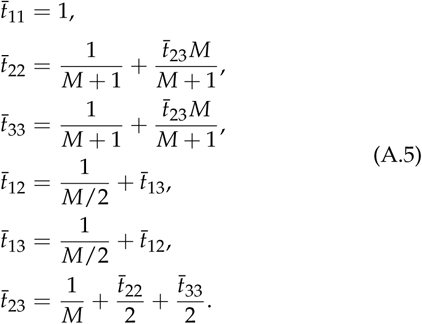

Note that this system is equivalent to considering that the isolated subpopulation 1 follows eq. A.2, that subpopulations 2 and 3 follow eq. A.3 with labels 2 and 3 in place of 1 and 2, and that coalescence times between subpopulations without migration (1 and 2 or 1 and 3) are infinite.

**Figure A1.**
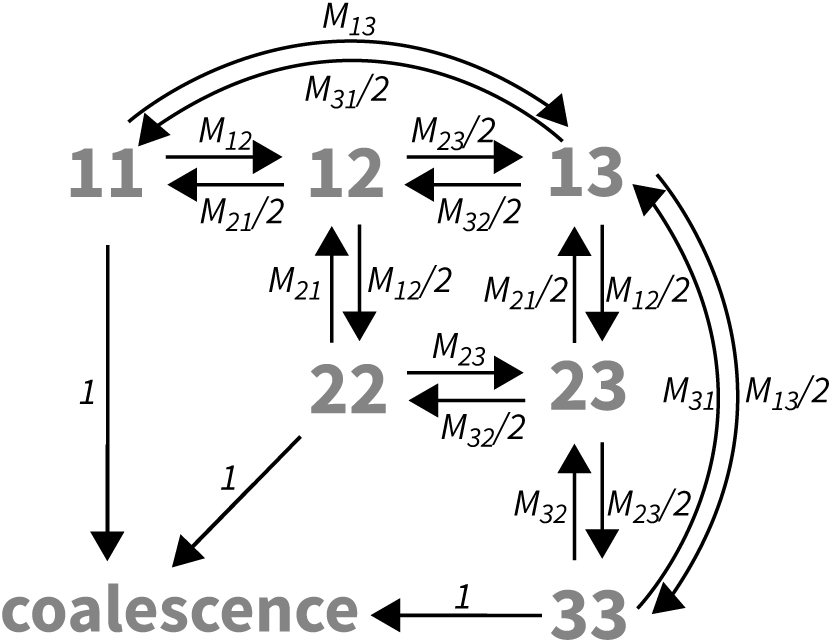
State diagram of the Markov chain representing the coalescent process of two lineages sampled in *K* = 3 subpopulations. States appear in gray and correspond to those presented in Figure 2; transition rates between states appear in black. *M*_*ij*_ corresponds to the scaled migration rate between subpopulations *i* and *j*. This diagram applies to all motifs with *K* = 3 subpopulations—motifs 4 to 6 in Figure 1. For example, motif 4 corresponds to the case where *M*_12_ = *M*_21_ = *M*_13_ = *M*_31_ = *M*_23_ = *M*_32_ = 0, and motif 5 corresponds to the case where *M*_12_ = *M*_21_ = *M* and *M*_13_ = *M*_31_ = *M*_23_ = *M*_32_ = 0.

We can solve this system using substitution, by first noting that *t*̄_22_ = *t*̄_33_, and by then substituting the expression for *t*̄_22_ into the equation of *t*̄_23_. We obtain (*t*̄_11_, *t*̄_22_, *t*̄_33_, *t*̄_12_, *t*̄_13_, *t*̄_23_) = (1,2,2, ∞, ∞, 2 + 1/*M*) as reported in Table 2.

### 4-vertex motifs

For four subpopulations, eq. A.1 simplifies to

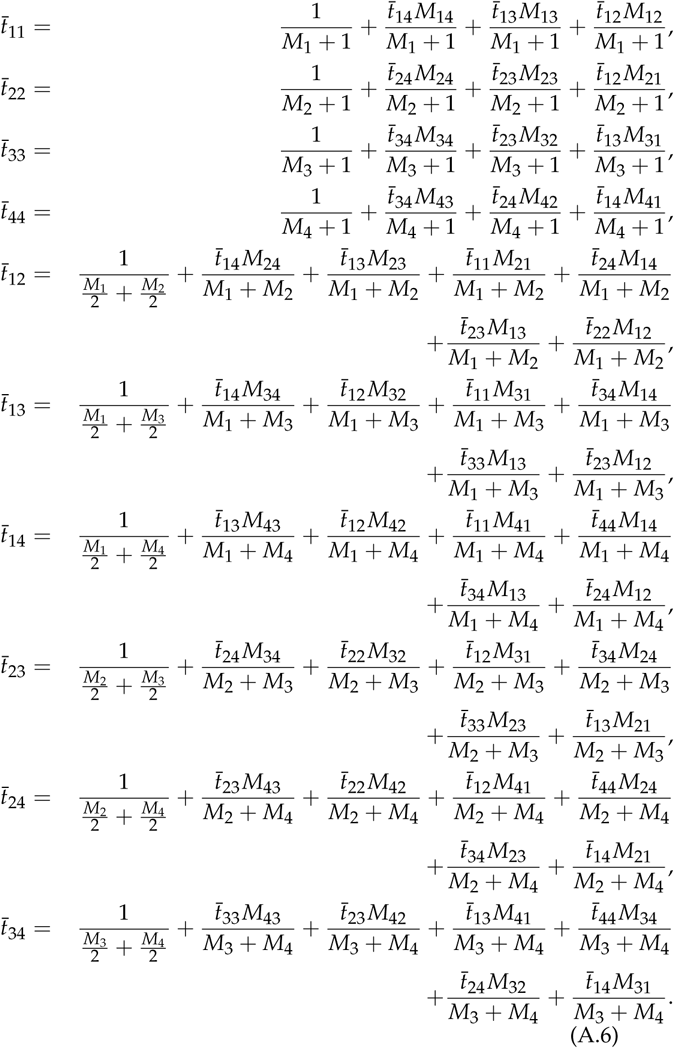

Similarly to the case of 3-vertex motifs, we set the values *M*_*ij*_ to reflect the network motifs of Figure 1, solve the system of equations using substitution or matrix inversion, and report the corresponding expected coalescence times in Table 3.

